# MTGCL: Multi-Task Graph Contrastive Learning for Identifying Cancer Driver Genes from Multi-omics Data

**DOI:** 10.1101/2023.10.13.562159

**Authors:** Ming-Yu Xie, Shao-Wu Zhang, Tong Zhang, Yan Li, Xiaodong Cui

**Author notes:** **Corresponding author.** Shao-Wu Zhang, School of Automation, Xiaodong Cui, School of Marine Science and Technology, Northwestern Polytechnical University, 127 West Youyi Road, Xi’an, Shaanxi, 710072, P.R.China. **Ming-Yu Xie** is currently working toward a PhD degree at the School of Automation from Northwestern Polytechnical University, China. He does researches in deep learning, machine learning and their applications to human cancer genomics. **Shao-Wu Zhang** received a PhD degree at the School of Automation from Northwestern Polytechnical University, China, at 2004. He has been a professor in Key Laboratory of Information Fusion Technology of Ministry of Education, School of Automation, Northwestern Polytechnical University, China. His current research interests include bioinformatics, complex networks and machine learning. **Tong Zhang** is a PhD candidate at the School of Automation from Northwestern Polytechnical University, China. He does researches in complex networks, machine learning and their applications to human cancer genomics. **Yan Li** is currently working toward a PhD degree at the School of Automation from Northwestern Polytechnical University, China. He does researches in complex networks and its applications to human cancer genomics. **Xiaodong Cui** received a PhD degree at the University of Texas at San Antonio, at 2016. He has been an associate professor in School of Marine Science and Technology, Northwestern Polytechnical University, China. His current research interests include bioinformatics, audio signal processing and machine learning.

## Abstract

Cancer is a complex disease that typically arises from the accumulation of mutations in driver genes. Identification of cancer driver genes is crucial for understanding the molecular mechanisms of cancer, and developing the targeted therapeutic approaches. With the development of high-throughput biological technology, a large amount of genomic data and protein interaction network data have been generated, which provides abundant data resources for identifying cancer driver genes through computational methods. Given the ability of graph neural networks to effectively integrate graph structure topology information and node features information, some graph neural network-based methods have been developed for identifying cancer driver genes. However, these methods suffer from the sparse supervised signals, and also neglect a large amount of unlabeled node information, thereby affecting their ability to identify cancer driver genes. To tackle these issues, in this work we propose a novel Multi-Task Graph Contrastive Learning framework (called MTGCL) to identify cancer driver genes. By using self-supervised graph contrastive learning to fully utilize the unlabeled node information, MTGCL designs an auxiliary task module to enhance the performance of the main task of driver gene identification. MTGCL simultaneously trains the auxiliary task and main task, and shares the graph convolutional encoder weights, so that the main task enhances the discriminative ability of the auxiliary task via supervised learning, whereas the auxiliary task exploits the unlabeled node information to refine the node representation learning of the main task. The experimental results on pan-cancer and some specific cancers demonstrate the effectiveness of MTGCL in identifying the cancer driver genes. In addition, integrating multi-omics features extracted from multiple cancer-related databases can greatly enhance the performance of identifying cancer driver genes, especially, somatic mutation features can effectively improve the performance of identifying specific cancer driver genes. The source code and data are available at https://github.com/NWPU-903PR/MTGCL.

**Author Summary:** Identifying cancer driver genes that causally contribute to cancer initiation and progression is essential for comprehending the molecular mechanisms of cancer and developing the targeted therapeutic strategies. However, wet-lab experiments are time-consuming and labor-intensive. The advent of high-throughput multi-omics technology provides an opportunity for identifying the cancer driver genes through data-driven computing approaches. Nevertheless, effectively integrating these omics data to identify cancer driver genes poses significant challenges. Existing computational methods exhibit certain limitations. For instance, conventional approaches (e.g., gene mutation frequency-based methods, network-based methods) often focus on a single omics data, while existing deep learning-based methods have not fully utilized the abundant unlabeled node information, so that their identification accuracy is not high enough. Thus, by fully utilizing multidimensional genomics data and molecular interaction networks, we propose a multi-task learning framework (called MTGCL) to identify cancer driver genes. MTGCL synergistically combines graph convolutional neural networks with graph contrastive learning. The experimental results validate the power of MTGCL for identifying cancer driver genes.

## Introduction

Cancer driver genes directly or indirectly participate in the formation and development of tumors, and their abnormal changes, such as gene mutations and copy number variations, can cause cell carcinogenesis, uncontrolled proliferation and metastasis of cancer cells, thus promoting tumor formation and progression[1–3]. Identifying cancer driver genes helps uncover the pathogenesis of cancer, develop more precise therapeutic schedule, and explore some new drug targets[4, 5]. Although wet-lab experiments can identify the cancer driver genes, it is time consuming and expensive. With the rapid development of biotechnology, and the implementation of projects such as The Cancer Genome Atlas (TCGA)[6] and the International Cancer Genome Consortium (ICGC)[7] and other projects[8, 9], a large amount of various omics data (e.g., mutation data, transcriptome data, proteome data, genome copy number change data) related to cancers have been generated. These omics data provide a guarantee for developing computational methods to identify cancer driver genes.

Existing computational methods of identifying cancer driver genes can be classified into three categories: gene mutation frequency-based methods, network-based methods, and machine learning-based methods. Gene mutation frequency-based methods identify the genes with higher mutation frequencies than the background mutations as cancer driver genes[10, 11]. These methods are limited by the selection of background mutation rate, making them difficult to distinguish high-frequency passenger genes and may miss some genes closely related to cancer but with lower mutation frequencies. Network-based methods assume that genes with important network structural properties[12, 13] or affecting the state of the entire molecular network[14, 15] are cancer driver genes. Due to the complex interrelationships among genes and the limited information on known gene relationships[16], network-based methods are limited by the incompleteness of the gene interaction network [17].

Machine learning-based methods of identifying cancer driver genes can be further divided into traditional machine learning methods and deep learning methods. Traditional machine learning methods usually select the effective features (e.g., gene mutation frequency, gene differential expression, and DNA methylation levels) to represent the genes, then feed these features into the classifier (e.g., support vector machine, random forest, and multilayer perceptron) to identify cancer driver genes[18–20]. Although these traditional machine learning-based methods have achieved better results in identifying cancer driver genes, they only utilize the genomic multi-omics features, which are insufficient to obtain the domain knowledge related to cancer. In recent years, deep learning models have made unprecedented progress in the fields of molecular biology and genomics. Some deep learning-based cancer driver gene identification methods have presented to identify cancer driver genes. Especially, graph neural networks that effectively integrate graph topology and node attribute information have been developed to identify cancer driver genes. For instance, EMOGI[21] primarily used graph convolutional networks (GCNs) to identify cancer driver genes by integrating protein-protein interaction networks and multiple genomics data (i.e., gene expression data, gene mutation frequency data, DNA methylation data, and copy number variation data), and also used the Layer-wise Relevance Propagation[22] to interpret and analyze the prediction results. MTGCN[23] employed a multi-task graph convolutional neural network to identify pan-cancer driver genes and specific cancer driver genes by optimizing both node prediction task and link prediction task during the learning process, and also concatenating the biological multi-omics features with network structure features obtained from DeepWalk[24]. Although EMOGI[21] and MTGCN[23] have achieved better results than the traditional machine learning-based cancer driver gene identification methods, they still have the following limitations: EMOGI only utilized a very sparse set of labeled genes information, which limits its performance improvement; MTGCN designed an link prediction auxiliary task to utilize the unlabeled gene information, but its prediction performance and computational efficiency is significantly affected by the sparsity and scale of protein-protein interaction (PPI) network.

In view of that self-supervised learning provides a feasible solution for graph representation learning by effectively exploiting the unlabeled data, especially graph contrastive learning (GCL) uses the data itself to provide supervised information to guide learning richer representation of each sample from a large amount of unlabeled data[25–27], in this work we introduce self-supervised graph contrastive learning to design an auxiliary task to share the graph convolutional encoder weights with the main task, and jointly train an end-to-end multi-task graph contrastive learning model to identify cancer driver genes. In sum, the contributions of this work are mainly as follows:

1. We introduce self-supervised graph contrastive learning to design an auxiliary task, and utilize the graph convolutional networks to plan the main task of node classification. The main task enhances the discriminative ability of the auxiliary task via supervised learning, while the auxiliary task refines the gene representation learning of the main task by exploiting the unlabeled node information.
2. We design a new convolutional layer structure for main task and auxiliary task, which can improve the performance of cancer driver gene identification by effectively integrating graph structure topology information and node features information.
3. We perform comprehensive experiments on pan-cancer and some specific cancers to demonstrate the effectiveness of MTGCL in identifying cancer driver genes.

## Results

### MTGCL architecture

MTGCL is an end-to-end framework (Fig 1) that effectively combines graph convolutional networks (GCNs) with graph contrastive learning (GCL) to identify cancer driver genes by using the molecular interaction networks and gene features as inputs. It generates scores for each gene, which represent the probability of the gene being a cancer driver gene. MTGCL mainly consists of a main task module and an auxiliary task module. The main task module is used to realize the semi-supervised node binary classification task using GCNs, while the auxiliary task is used to implement the node representation learning task using graph contrastive learning. We simultaneously train the two task modules and let them share the graph convolutional encoder weights, so that the main task enhances the discriminative ability of the auxiliary task via supervised learning, whereas the auxiliary task exploits the unlabeled node information to refine the node representation learning, thereby improving the classification performance of the main task. This learning paradigm enables MTGCL to learn more cancer-related representations for each gene.

**Fig 1.**
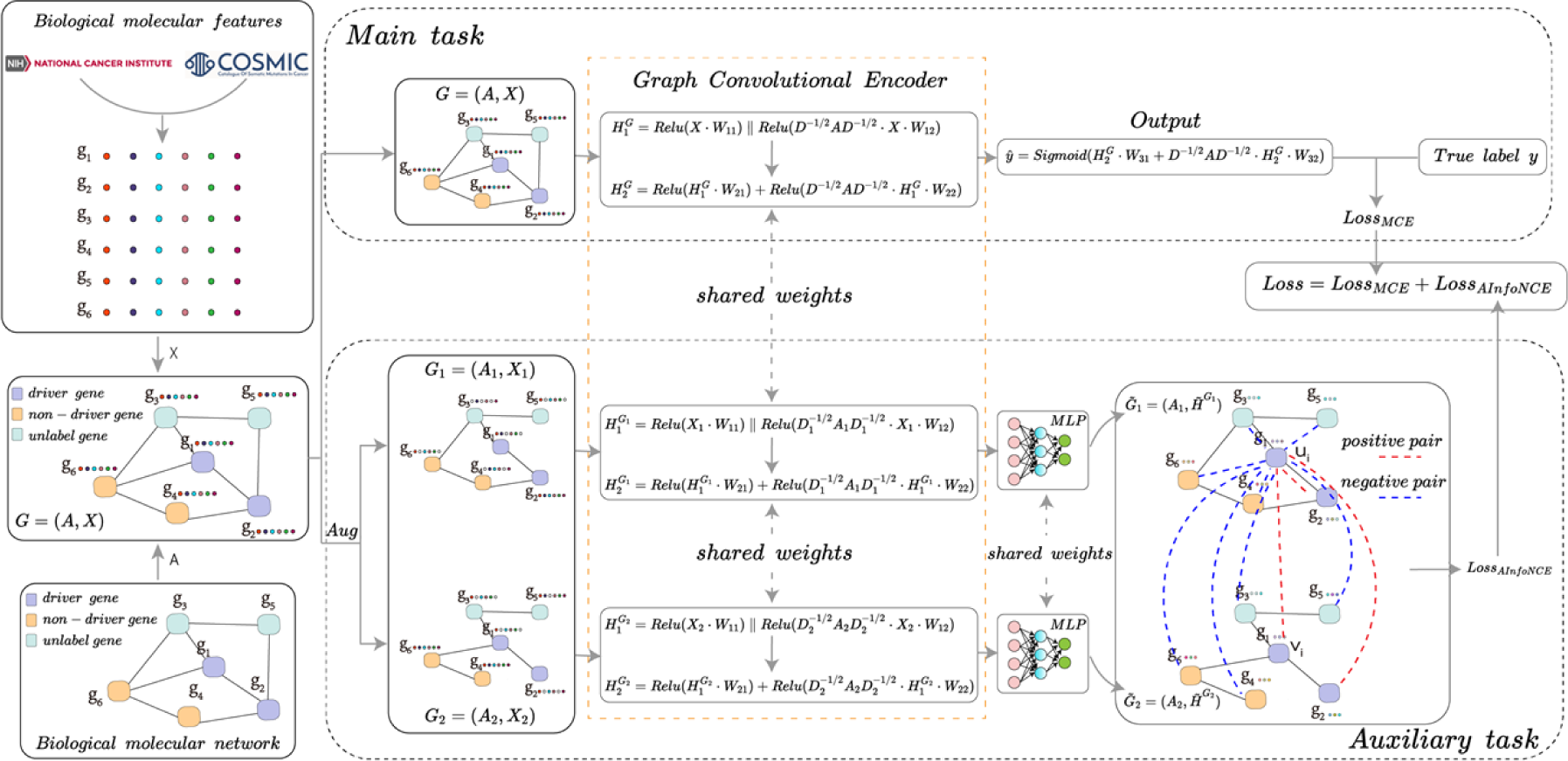
Architecture of MTGCL. MTGCL mainly consists of two task modules: the main task module that implements the semi-supervised graph convolutional neural network node classification task, and the auxiliary task module that implements the graph contrastive learning node feature representation learning task. These two modules are trained simultaneously by sharing the weights in the graph convolutional encoder. The inputs of MTGCL consist of the adjacency matrix *A* and the node feature matrix *X*. The output 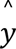 is a vector in which each element represents the probability of each gene being predicted as a cancer driver gene. ‘Aug’ means graph augmentation operation. ′||′ indicates concatenation operation.

### Performance of MTGCL and other seven contrastive methods for identifying pan-cancer driver genes

In order to evaluate the performance of MTGCL for identifying pan-cancer driver genes, we compared our MTGCL with other seven methods on CPDB, STRING and PathNet networks. Seven contrastive methods are the five baseline methods of Multi-Layer Perceptron (MLP), DeepWalk[24], GCN[28], GAT[29], ChebConv[30], and two state-of-the-art methods of EMOGI[21] and MTGCN[23]. The hyperparameters for each method can be found in the Supplementary A. The metrics of the area under the precision-recall curve (AUPR) and the area under the Receiver Operating Characteristic curve (AUC) are used to evaluate the performance of MTGCL and other seven comparative methods. AUPR punishes much more the existence of false-positive cancer driver genes among the best-ranked prediction scores, therefore it is a more significant quality metric than AUC in the class-imbalanced task[31]. The results of MTGCL and other seven comparative methods in 5-fold cross-validation (5CV) test are shown in Table 1. We also performed the *t* hypothesis test (*t*-test) to verify whether the performance improvement of MTGCL over the other contrastive methods is significant. Considering that ChebConv, EMOGI and MTGCL have better results than other four contrastive methods, we just provide the significance test results (S1 Fig in supplementary B) of MTGCL compared to ChebConv, EMOGI and MTGCN methods on CPDB, STRING and PathNet networks.

**Table 1.** Performance of MTGCL and other contrastive methods on CPDB, STRING and PathNet networks in 5CVtest.

|  | CPDB |  | STRING |  | PathNet |  |
| --- | --- | --- | --- | --- | --- | --- |
|  | AUPR | AUC | AUPR | AUC | AUPR | AUC |
| MLP | 0.6706±0.0015 | 0.8136±0.0012 | 0.6782±0.0029 | 0.8464±0.0007 | 0.7960±0.0033 | 0.8062±0.0027 |
| DeepWalk | 0.7559±0.0023 | 0.8735±0.0008 | 0.7280±0.0037 | 0.8697±0.0012 | 0.7008±0.0065 | 0.7336±0.0053 |
| GCN | 0.7319±0.0030 | 0.8682±0.0015 | 0.7086±0.0026 | 0.8754±0.0013 | 0.7367±0.0064 | 0.7587±0.0059 |
| GAT | 0.7025±0.0261 | 0.8577±0.0075 | 0.7176±0.0141 | 0.8780±0.0044 | 0.7378±0.0122 | 0.7885±0.0057 |
| ChebConv | 0.8202±0.0028 | 0.9022±0.0015 | 0.8101±0.0027 | 0.9125±0.0011 | 0.8203±0.0038 | 0.8352±0.0023 |
| EMOGI | 0.8107±0.0030 | 0.8984±0.0020 | 0.8038±0.0039 | 0.9106±0.0016 | 0.8089±0.0051 | 0.8267±0.0028 |
| MTGCN | 0.8265±0.0016 | 0.9053±0.0014 | 0.8093±0.0037 | 0.9130±0.0016 | 0.7932±0.0049 | 0.8112±0.0032 |
| MTGCL | **0.8316±0.0022** | **0.9084±0.0011** | **0.8240±0.0027** | **0.9206±0.0013** | **0.8531±0.0028** | **0.8622±0.0032** |

From Table 1 and S1 Fig, we can see that the performance of our MTGCL outperforms all of the other seven methods of MLP, DeepWalk, GCN, GAT, ChebConv EMOGI and MTGCN for identifying the cancer diver genes on three molecular networks of CPDB, STRING, and PathNet. The AUPR and AUC values of MTGCL on the CPDB network are 0.8316 and 0.9084, which are 0.0051∼0.1610 and 0.0031∼0.0948 higher than those of the other seven methods, respectively; AUPRC and AUC values of MTGCL on STRING network are 0.8240 and 0.9206, which are 0.0147∼0.1458 and 0.0076∼0.0742 higher than those of other seven methods, respectively; AUPRC and AUC values of MTGCL on PathNet network are 0.8531 and 0.8622, which are 0.0328–0.1523 and 0.027–0.1286 higher than those of other seven methods, respectively. Moreover, the AUPR performance improvement of MTGCL is statistically significant compared to these methods with similar performance, including MTGCN, EMOGI, and ChebConv. These results demonstrate that our MTGCL method has superior performance in identifying the pan-cancer driver genes.

In addition, we also compared the running time and convergence speed of MTGCL with MTGCN that also uses the GCN multi-task learning strategy (i.e., taking the node prediction as the main task and the link prediction as the auxiliary task) under the same experimental environment. On CPDB network, MTGCL takes 203 seconds to run 1000 epochs, while MTGCN takes 1,004 seconds; the convergence iteration number of MTGCL is 1,900, while MTGCN is 2,500. On STRING network, MTGCL takes 185 seconds to run 1000 epochs, while MTGCN takes 1,368 seconds; the convergence iteration number of MTGCL is 1,900, while MTGCN is 2,800. On PathNet network, MTGCL takes 89 seconds to run 1000 epochs, while MTGCN takes 342 seconds; the convergence iteration number of MTGCL is 700, while MTGCN is 800. Above results demonstrate that our MTGCL not only performs better than MTGCN in identifying pan-cancer driver genes, but also has higher computational efficiency than MTGCN.

### Ablation experiments of diverse architecture components in MTGCL

To evaluate the contributions of diverse architecture components in our MTGCL, we conducted ablation experiments on CPDB, STRING and PathNet networks in 5CV test, and also analyzed the impact of omics features from two different databases (i.e., TCGA and COSMIC) on MTGCL. The ablation experimental results of MTGCL are shown in Table 2. In Table 2, MTGCL_-GCL_ denotes that we remove the contrastive learning module from MTGCL. MTGCN_^ChebConv_ denotes that we use Chebyshev convolutional layer instead of our designed graph convolutional encoder in MTGCL. MTGCL_XT_ denotes that we only use the 48-dimensional features extracted from TCGA in in MTGCL. MTGCL_XC_ denotes that we only use the 37-dimensional features extracted from COSMIC in MTGCL. We also performed the *t*-test to verify whether the contributions of diverse architectural components in MTGCL are statistically significant. S2 Fig (in supplementary B) is the AUPR statistical significance test results of MTGCL compared to its variations (i.e., MTGCL_-GCL_, MTGCN_^ChebConv_, MTGCL_XT_ and MTGCL_XC_).

**Table 2.** Ablation experiment results of MTGCL on CPDB, STRING and PathNet networks.

|  | CPDB |  | STRING |  | PathNet |  |
| --- | --- | --- | --- | --- | --- | --- |
|  | AUPR | AUC | AUPR | AUC | AUPR | AUC |
| $\text{MTGCL}_{\text{-GCL}}$ | 0.8252±0.0030 | 0.9046±0.0016 | 0.8137±0.0030 | 0.9156±0.0012 | 0.8498±0.0033 | 0.8569±0.0032 |
| $\text{MTGCN}^{\wedge}_{\text{ChebConv}}$ | 0.8255±0.0017 | 0.9063±0.0008 | 0.8185±0.0018 | 0.9173±0.0006 | 0.8244±0.0054 | 0.8386±0.0030 |
| $\text{MTGCL}_{\text{-XT}}$ | 0.8165±0.0035 | 0.9005±0.0011 | 0.7976±0.0015 | 0.9071±0.0010 | 0.8323±0.0029 | 0.8453±0.0025 |
| $\text{MTGCL}_{\text{-XC}}$ | 0.8046±0.0024 | 0.8918±0.0012 | 0.8091±0.0037 | 0.9143±0.0015 | 0.8451±0.0045 | 0.8538±0.0042 |
| **MTGCL** | **0.8316±0.0022** | **0.9084±0.0011** | **0.8240±0.0027** | **0.9206±0.0013** | **0.8531±0.0028** | **0.8622±0.0032** |

As shown in Table 2 and S2 Fig, we can see that the AUPR values of MTGCL on the CPDB, STRING and PathNet networks are 0.8316, 0.8240, and 0.8513, which are 0.0064, 0.0103 and 0.0033 higher than that of MTGCL_-GCL_, respectively, and the AUPR performance improvement of MTGCL is statistically significant compared to MTGCL_-GCL_, indicating that introducing graph contrastive learning to design the auxiliary task can improve the performance of MTGCL in identifying the pan-cancer driver genes. The AUPR values of MTGCL on the CPDB, STRING and PathNet networks are 0.8316, 0.8240, and 0.8513, which are 0.0061, 0.0055, and 0.0287 higher than that of MTGCN_^ChebConv_, respectively, our designed convolutional layer can enhance the performance of MTGCL than the Chebyshev convolutional layer. The AUPR value of MTGCL on the PathNet network are 0.8513, which is 0.0208, 0.008 higher than that of MTGCL_XT_ and MTGCL_XC_, respectively, indicating that the integration of omics features extracted from TCGA and COSMIC databases can improve the performance of MTGCL. The results in Table 2 show that all the proposed components for building MTGCL are valid and contribute to the final performance of MTGCL, and integrating omics features extracted from multiple cancer databases can effectively improve the performance of MTGCL for identifying pan-cancer driver genes.

### Performance analysis of MTGCL in identifying novel pan-cancer driver genes

In this section, we analyze the performance of MTGCL in identifying novel pan-cancer driver genes on three molecular networks (i.e., CPDB, STRING and PathNet networks). For each molecular network, we run MTGCL 10 times in 5CV test to calculate the average prediction score of genes as the final prediction score for the following analysis. We first analyze whether there is a significant difference in prediction scores between known cancer driver genes (KCGs) and candidate cancer genes (CCGs)/ non-cancer genes (noCGs)/ other genes, as well as between CCGs and noCGs/ other genes. Fig 2a presents the prediction scores of KCGs, CCGs, noCGs and other genes on three molecular networks, and also provides the statistical significance *t*-test results between KCGs and CCGs/ noCGs/ other genes, as well as between CCGs and noCGs/ other genes. As shown in Fig 2a, we can see that on three molecular networks, the scores of KCGs are significantly higher than that of CCGs, noCGs and other genes, and the scores of CCGs are also significantly higher than that of noCGs and other genes, indicating that MTGCL can not only accurately identify KCGs, but also predict CCGs that potentially driver cancer development.

**Fig 2.**
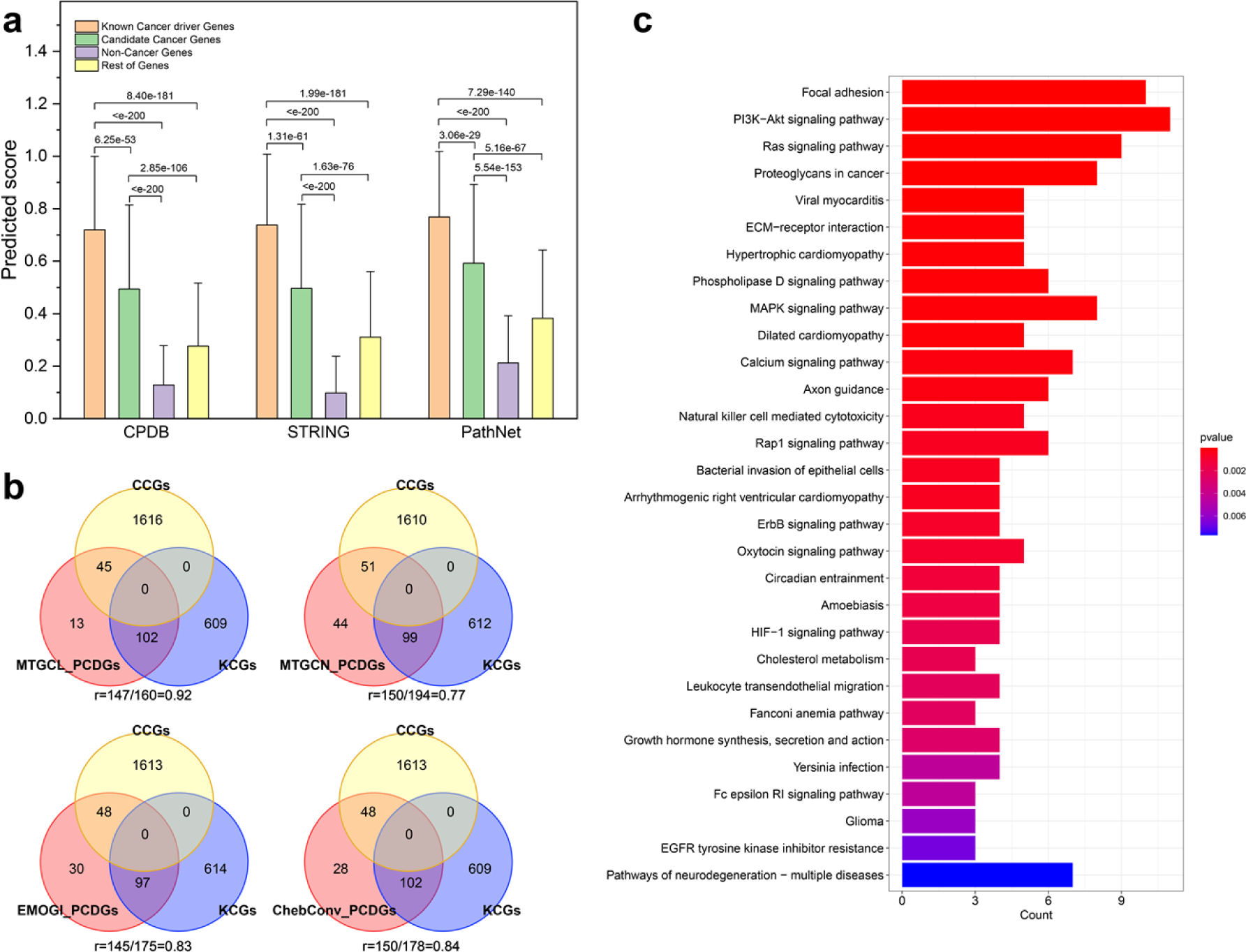
Performance of MTGCL for identifying novel pan-cancer candidate driver genes on CPDB, STRING and PathNet networks. (a) The average scores of KCGs, CCGs, noCGs, and other genes predicted by MTGCL on CPDB, STRING and PathNet networks, respectively. (b) Venn diagram of PCDGs, KCGs and CCGs obtained from MTGCL, MTGCN, EMOGI and ChebConv methods. (c) Results of KEGG pathway enrichment analysis of 57 NPCGs identified by MTGCL. The horizontal axis represents the number of NPCGs involved in the corresponding pathway. The color bar denotes the *p*-value.

Secondly, we select the top 100 genes with the highest prediction scores from CPDB, STRING, and PathNet networks, respectively. These genes are merged into a set (a total of 160 genes in this set), which are used as the Predicted Cancer Driver Genes (PCDGs) of MTGCL. Using the same strategy as above, we can obtain the PCDGs sets of MTGCN, EMOGI, and ChebConv, which contain 194, 175, and 178 genes, respectively. The Venn diagram in Fig 2b presents the number of gene overlaps among PCDGs, KCGs and CCGs, as well as the overlap rate between PCDGs and KCGs+CCGs for MTGCL, MTGCN, EMOGI and ChebConv methods. From Fig 2b, we can see that the overlap rate (0.92) of MTGCL is higher than that of MTGCN (0.77), EMOGI (0.83) and ChebConv (0.84), indicating the prediction results of our MTGCL have higher confidence level.

Thirdly, we remove 103 genes that are already part of the training or test set in three networks from PCDGs set of MTGCL, and treat the reminding genes as newly predicted cancer driver genes (named NPCGs). As a result, we obtained 57 NPCGs (S1 Table in supplementary B), of which 44 genes are labeled as CCGs, 8 genes (i.e., *FN1, PRKCA, APP, SHC1, TCF4, TEK, LAMA1*, and *GEN1*) are annotated as driver genes, or oncogenes/tumor suppressor genes in the CancerMine database[32], and 4 genes (i.e., *FRAS1, RYR3, SLX4,* and *RMI1*) are the essential genes in Project Achilles[33]. Out of the 57 NPCGs, 56 genes have at least one evidence supporting them the potential as cancer driver genes, while another gene (*SLX1A*) encodes a protein that regulates genomic stability.

Finally, we perform KEGG pathway enrichment analysis on the 57 NPCGs identified by MTGCL. Fig 2c shows the top thirty enriched pathways (the detailed KEGG pathway enrichment analysis results of 57 NPCGs are presented in S4 Table (in Supplementary C)), we can see that NPCGs are enriched into many KEGG pathways related to cancer, such as Focal adhesion, PI3K-Akt signaling pathway, Ras signaling pathway, Proteoglycans in cancer, MAPK signaling pathway, etc. And many genes in NPCGs play important roles in these cancer-related pathways. For example, *PTK2* (also known as Focal Adhesion Kinase, *FAK*) is a key gene encoding the focal adhesion kinase, a critical protein in the focal adhesion pathway. Previous studies have shown that *FAK* plays a role in breast tumor progression, intestinal tumorigenesis, and squamous cell carcinoma [34–36]. The *FN1* gene is important in both the Focal Adhesion and PI3K-Akt signaling pathways. More studies have demonstrated that *FN1* is highly expressed in breast cancer [37, 38] and involves in the interaction between breast cancer cells and immune cells in the tumor microenvironment [39].

### Performance contrastive of MTGCL with other seven methods for identifying specific cancer driver genes

To validate the effectiveness of MTGCL in identifying specific cancer driver genes, we perform MTGCL and other seven methods on CPDB network to identify the driver genes of Breast Cancer (BRCA) and Liver Hepatocellular Carcinoma (LIHC), respectively. We just use the breast-related data to extract the gene node features, and liver-related data to extract the gene node features in the CPDB network. That is, we extracted 40-dimensional features (*X*_40_) for each gene, which consist of 3-dimensional features (*X*_3_) of single nucleotide mutation frequency, gene differential methylation, and gene differential expression obtained from TCGA, as well as 37-dimensional somatic mutation features (*X*_37_) collected from the COSMIC database. The genes annotated in the NCG database[9] that are associated with BRCA/LIHC cancer are used as the positive samples. After mapping them to the CPDB network, we can obtain 201 positive samples for BRCA and 82 positive samples for LIHC. The 2,187 negative sample genes determined in pan-cancer driver gene identification are taken as the negative samples for identifying BRCA/LIHC cancer driver genes. The optimal hyperparameters used in MTGCL and other seven methods can be found in “Hyperparameter settings” in Supplementary A.

Table 3 presents the AUPR and AUC values of MTGCL and other seven methods for identifying BRCA and LIHC cancer driver genes on CPDB network in 5CV test. S3 Table (in Supplementary B) gives the significance *t*-test results of MTGCL compared to other methods on BRCA and LIHC cancers. From Table 3 and S3 Table, we can see that the performance of our MTGCL is still superior to other seven methods in identifying BRCA and LIHC cancer driver genes, indicating that our MTGCL can also be effectively used to identify the specific cancer genes. In addition, we also investigated the impact of *X*_3_ and *X*_37_ features on MTGCL (S3 Fig), finding that somatic mutation features (*X*_37_) has a significant contribution to improve the performance of MTGCL compared to the features (*X*_3_) of single nucleotide mutation frequency, gene differential methylation, and gene differential expression. However, EMOGI[21] and MTGCN[23] only use *X*_3_features to identify the BRCA/LIHC driver genes in their published original literatures.

**Table 3.** Performance of MTGCL and other seven methods on BRCA and LIHC.

|  | BRCA |  | LIHC |  |
| --- | --- | --- | --- | --- |
|  | AUPR | AUC | AUPR | AUC |
| MLP | 0.5167±0.0162 | 0.8797±0.0065 | 0.5745±0.0089 | 0.8735±0.0049 |
| DeepWalk | 0.6376±0.0041 | 0.8975±0.0011 | 0.4036±0.0081 | 0.8732±0.0033 |
| GCN | 0.5723±0.0110 | 0.8952±0.0031 | 0.2943±0.0735 | 0.8207±0.0241 |
| GAT | 0.5149±0.0560 | 0.8665±0.0291 | 0.3783±0.0633 | 0.8602±0.0145 |
| ChebConv | 0.7716±0.0052 | 0.9373±0.0029 | 0.6337±0.0112 | 0.9005±0.0052 |
| EMOGI | 0.7568±0.0108 | 0.9306±0.0043 | 0.6043±0.0159 | 0.8884±0.0051 |
| MTGCN | 0.7723±0.0072 | 0.9415±0.0033 | 0.6439±0.0147 | 0.9007±0.0049 |
| MTGCL | **0.8034±0.0041** | **0.9484±0.0022** | **0.6751±0.0135** | **0.9128±0.0040** |

## Discussion

In this work, we developed a novel framework of MTGCL to identify the cancer driver genes. MTGCL is a multi-task learning model framework that introduces the graph contrastive learning to design the auxiliary task and proposes a novel graph convolutional layer structure for the main task of GCN-based node classification. The auxiliary task utilizes a large amount of unlabeled node information to boost the node representation learning of the main task, thereby improving the performance of cancer driver gene identification. The main task enhances the discriminative ability of the auxiliary task via supervised learning. MTGCL jointly trains the auxiliary and main tasks by sharing the propagation layer parameters to improve the model performance and reduce computational costs. The results on three molecular networks of GPDB, STRING and PathNet show that MTGCL outperforms other existing state-of-the-art methods in identifying pan-cancer and specific cancer driver genes. The ablation experimental results demonstrate that our proposed new graph convolution strategy can effectively improve the performance of identifying cancer driver genes, and the graph contrastive learning auxiliary task can boost the main task gene representation learning for further improving model performance. In addition, integrating omics features extracted from multiple cancer-related databases can greatly enhance the performance of identifying cancer driver genes. Especially, somatic mutation features can effectively improve the performance of identifying specific cancer driver genes. The analysis results of top 100 predicted cancer driver genes demonstrate that the confidence level of our MTGCL prediction results is relatively high.

Although MTGCL has achieved good performance in identifying pan-cancer and some specific cancers (i.e., BRCA and LIHC) driver genes, it can be improved from the following four aspects. Firstly, MTGCL should consider introducing more biological prior information, developing efficient feature extraction and integration approaches to effectively represent genes, even though it considers more omics data related to gene mutation types, achieving a significant performance improvement of cancer driver gene identification. Secondly, different molecular networks have a certain impact on the identification results of MTGCL, we should consider fully utilizing more known molecular interaction information to build a more complete network. Thirdly, MTGCL introduces graph contrastive learning to design an auxiliary task to alleviate the limited label problem, but it still cannot completely get rid of its dependence on labels, therefore we should develop an effective self-supervised learning method to reduce dependence on label information.

Our MTGCL is a very flexible framework, especially in the design of the convolutional layer. Here, we chose to use the first-order neighbor nodes (not including the node itself) degree normalized average for neighbor aggregation, which is very suitable for the data characteristics of the problem studied in this work. However, in other fields, this aggregation method can be replaced by other methods such as graph attention mechanism, etc.

## Materials and Methods

### An overview of the data

MTGCL utilizes the multi-omics data and three different biological molecular networks to identify cancer driver genes. The multi-omics data are used to build the gene feature matrix *X* ∈ *R*^*N*×85^, which contains *N* genes, and each gene is represented by an 85-dimensional feature vector *x*_*i*_ ∈ *R*^1×85^. The 85 features in *x*_*i*_ are composed of two parts: 1) 48 features were obtained from the gene expression data, gene mutation data, and DNA methylation data of 16 different cancer types from The Cancer Genome Atlas (TCGA) database[6] by performing the same data preprocessing pipeline as EMOGI; 2) 37 features were related to different mutation types, which are obtained from somatic mutation data of 37 primary tissues in COSMIC[8] database by performing the same data collection and preprocessing pipeline as cTaG[40].

Three biological molecular networks are CPDB network, STRING network and PathNet network. CPDB network is a human functional interaction network that includes human binary and complex protein-protein, genetic, metabolic, signaling, gene regulatory and drug-target interactions, as well as biochemical pathways, and it is derived from the Consensus Pathway database[41]. STRING network is a protein-protein association network, which is derived from the STRING database[42]. PathNet network is the pathway network that includes KEGG and Reactome pathways[43]. The detailed processing procedures of multi-omics data and three molecular networks are provided in Supplementary A.

For identifying pan-cancer driver cancers, we choose the positive and negative samples with the same strategy as EMOGI. The positive samples (i.e., cancer driver genes) are the union of the known cancer driver genes (KCGs) obtained from the expert-curated list in The Network of Cancer Genes (NCG) [9], the Cancer Gene Census (CGCs) genes from the COSMIC[8] database, and high-confidence genes mined from literature [44]. The negative samples(i.e., non-cancer driver genes) are obtained by iteratively removing genes from the positive samples, NCG[9] candidate cancer driver genes, the genes associated with ‘pathways in cancer’ in the KEGG database[45], the disease genes in OMIM[46], the genes predicted to be involved with cancer by MSigDB[47], and the genes related to the expression of cancer genes[48]. Statistical results of CPDB, STRING and PathNet networks are shown in S2 Table (in Supplementary B).

### Main task module of MTGCL

In this section, we first analyzed the performance differences of using different graph convolutional structure in the main task module in which it adopts semi-supervised GCN model to achieve node classification task (i.e., identifying cancer drive genes), especially analyzing the reasons for performance improvement of ChebConv[30] (used in EMOGI[21] and MTGCN[23]) compared to GCN[28] and GAT[29]. Then, we presented a new graph convolutional layer structure for the main task and auxiliary task.

From the comparative experimental results (R1 Fig in Supplementary A) of using different graph convolutional structures, we can find that ChebConv can better integrate the graph structure topology information and omics feature information of nodes than GCN and GAT, thereby improving the performance of identifying cancer driver genes. This is because ChebConv performs message aggregation by separating the mapping matrices of node self-features (ego-features) from its neighboring features aggregated (neighborhood-features), while GCN and GAT share the mapping matrix. Therefore, ChebConv can avoid the discriminating features of a driver gene being smoothed during message aggregation, thereby alleviating the over-smoothing problem in GCN and GAT. That is, separating the feature embedding matrices for the ego-features and neighborhood-features during each convolution operation can enhance the model’s utilization of node feature information, thereby improving the model’s performance in identifying cancer driver genes. Due to the fact that ChebConv is a frequency-based graph convolution structure, it is difficult to understand and lacks flexibility in its convolutional structure. Therefore, leveraging the aforementioned discovered characteristic, we designed a new convolutional layer structure (Equation 1) from the perspective of the spatial domain.

For network *G*(*E*, *A*, *V*, *X*) = *G*(*A*, *X*), where *E* is the edge set of molecular network, *A* ∈ *R*^*N*×*N*^ is the adjacency matrix, *V* is the node set, *N* = |*V*| is the number of nodes in the network, *X* is the matrix consisting of the content features associated with each molecular node, the representation of our proposed graph convolutional layer structure at the *k*-th layer can be formulated as:

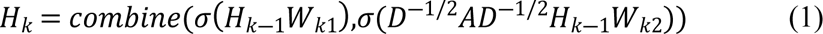

where, *H*_0_ = X, *D* is the degree matrix (*D_ii_* = ∑ *_j_ A_ij_*), *σ*(·) is the nonlinearity activation function (*i.e.*, ReLU, Sigmoid). *W*_*k*1_and *W*_*k*2_ represent the learnable parameter matrices of the ego-features and neighborhood-features of the node at the (*k*-1)-th layer, respectively. Operator *combine*(*P*, *Q*) represents the operation of summing (or concatenating, etc.) *P* and *Q*. In this work, we choose the concatenation operation in the first layer, because the first-order neighbors of a node have greatest impact on that node, and evaluating the relationship between the node own features and its first-order neighbor features is more conducive to learning more reliable representations. For the second layer and subsequent layers, we choose the summation operation, because their outputs need to be input into the auxiliary task of graph contrastive learning, and too high dimension of node features will increase the computational burden. For the second layer and subsequent layers, we choose the summation operation, because the output of the last layer of the graph convolutional encoder need to be input into the auxiliary task of graph contrastive learning, and too high dimension of node features will increase the computational burden. Considering our computer configuration and the model performance improvement, here we set *k* = 2. The output 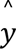 of the main task can be formulated as:

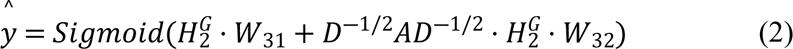

where, *W*_31_,*W*_32_ ∈ *R*^*N*∗1^represent the learnable parameter matrices of the ego-features and neighborhood-features of the node at the 3-th layer (i.e., output layer), respectively.

In the training process of MTGCL, we adopt following cross-entropy loss function *Loss*_*MCE*_ to optimize the main task.

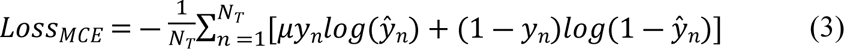

where, 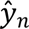 is the predicted score of *n*-th gene, *y*_*n*_ is the true label of *n*-th gene, *N*_*T*_ is the total number of gene in the training set, μ is a balance factor that controls the weight of positive samples, here we set μ to 1.

### Auxiliary task module of MTGCL

Due to the sparsity of known cancer driver genes, the prediction performance will be limited if only adopting the supervised learning models to identify cancer driver gene. In view of that graph contrastive learning can utilize the data itself to guide the learning of effective representations for each node from a large amount of the unlabeled node data[25–27]. In this work, we introduce the graph contrastive learning to design the auxiliary task module to refine the node representation learning of main task module by exploiting the unlabeled node information.

First, we generate two graph views (i.e., *G*_1_ = (*A*_1_,*X*_1_) and *G*_2_ = (*A*_2_,*X*_2_)) by performing stochastic graph augmentation on the network *G* = (*A*,*X*). That is, a certain proportion of edges in *G* = (*A*,*X*) are randomly deleted and a certain proportion of feature values are masked (i.e., set to 0) to generate two different graph views *G*_1_ = (*A*_1_,*X*_1_) and *G*_2_ = (*A*_2_,*X*_2_), respectively. Then, we input *G*_1_ = (*A*_1_,*X*_1_) and *G*_2_ = (*A*_2_,*X*_2_) into a pair of the shared graph convolutional encoders to extract two node representation matrices 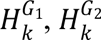.

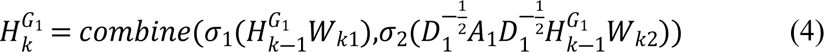

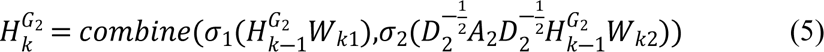

where, *D*_1_ is the degree matrix of the augmented graph *G*_1_(*A*_1_,*X*_1_), *D*_2_ is the degree matrix of the augmented graph *G*_2_(*A*_2_,*X*_2_).

After that, we utilize a pair of the shared multi-layer perceptron (MLP) to project the node representation from both views (*i.e.*, *G*_1_ and *G*_2_) into the mapping space to generate the mapping node representation matrices 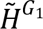 and 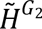, which are used to construct graphs 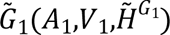 and 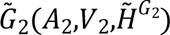 for node-level graph contrastive learning, where *V*_1_and *V*_2_are the node sets of graphs 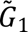 and 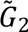, respectively. Lastly, we use the semi-supervised contrastive learning objective[49, 50] to optimize the auxiliary task, which allows the nodes of the same type to cluster together and separates the nodes of different types, while retaining a large amount of information from unlabeled nodes.

Let *u*_*i*_ ∈ V_1_ represents the *i*-th node in view 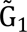 and *v*_*j*_ ∈ V_2_ represents the *j*-th node in view 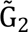, and the nodes in view 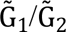 are divided into three sets: the training positive sample node set 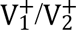, the training negative sample node set 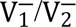, and the remaining sample node set 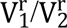. For the training sample nodes, we adopt the supervised contrastive learning to design the loss function 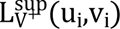 for each positive pair (*u*,*v*), and 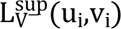 for each negative pair (u,v). For the remaining sample nodes, we employ the self-supervised contrastive learning to design the loss function 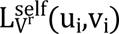 for each pair (u_i_,v_i_).

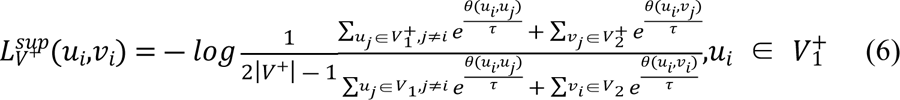

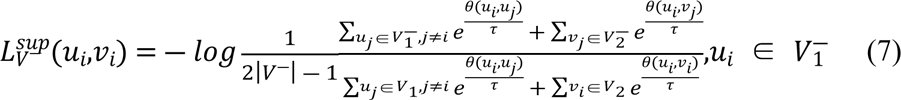

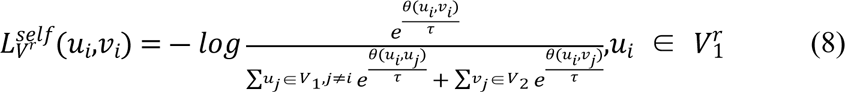

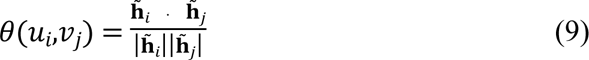

where 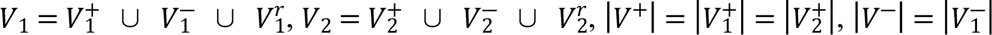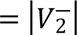. τ is a temperature parameter. 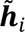 is the representation vector of *i*-th node in 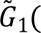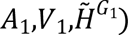 and 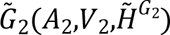, which is used to build matrix 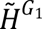 and 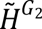. θ(*u*_*i*_,*v*_*j*_) is the cosine value used to measure the similarity of representations two nodes *u*_*i*_and *v*_*j*_.

Since two views are symmetric, the loss function for another view is similarly defined as *L*(*v*_*i*_,*u*_*i*_). Therefore, we define the following loss function *LOSS*_*AInfoNCE*_ to optimize the auxiliary task.

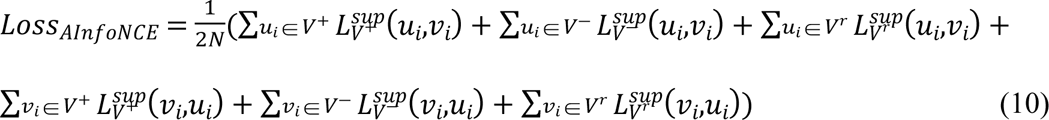

Taking into account both the main task loss and the auxiliary task loss, the total loss function of MTGCL is defined as follows:

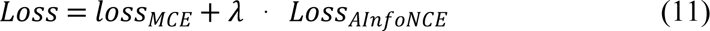

where λ is a factor coefficient, which are used to adjust the role of the auxiliary task in the total loss of MTGCL.

## Supporting information

**Supplementary A.** This supplementary materials include the following contexts: the preprocessing process of molecular omics data and molecular networks; analyzing the performance differences in cancer driver gene identification task with different convolutional layer structures; the hyperparameter settings of MTGCL and other comparative methods.

**Supplementary B.** It provides some supplementary Figures (S1 Fig - S3 Fig) and Tables (S1 Table - S3Table).

**S1 Fig. The AUPR statistical significance test results of MTGCL compared to other three methods of ChebConv, EMOGI and MTGCL on CPDB, STRING and PathNet networks.**

**S2 Fig. The AUPR statistical significance test results of MTGCL compared to its variations in ablation experiments.**

**S3 Fig. Results of MTGCL using different omics features for identifying BRCA and LIHC cancer driver genes.**

**S1 Table. List of the newly predicted cancer driver genes.**

**S2 Table. Statistical results of CPDB, STRING and PathNet networks.**

**S3 Table. Significance t-test results of MTGCL compared to other methods on BRCA and LIHC cancers.**

**Supplementary C.**

**S4 Table: Details of the KEGG enrichment analysis results for the 57 NPCGs by MTGCL.**

## Funding

This work was supported by the National Natural Science Foundation of China (Nos. 62173271 and 61873202) awarded to SWZ; the National Natural Science Foundation of China (No.62003273) awarded to XC; and was also partially supported by the Natural Science Basic Research Program of Shaanxi (No. 2020JQ-217) awarded to XC. The funders had no role in study design, data collection and analysis, decision to publish, or preparation of the manuscript.

## Author Contributions

**Conceptualization:** MYX SWZ XC.

**Data curation:** MYX TZ.

**Formal analysis:** MYX.

**Funding acquisition:** MYX SWZ XC.

**Investigation:** MYX YL.

**Methodology:** MYX.

**Project administration:** SWZ.

**Resources:** MYX YL.

**Software:** MYX.

**Supervision:** SWZ XC.

**Validation:** MYX YL.

**Visualization:** MYX.

**Writing – original draft:** MYX.

**Writing – review & editing:** SWZ XC TZ YL.

## Notes

### Competing Interest Statement

The authors have declared no competing interest.

